# Fractionated proton and photon FLASH irradiation mitigates radiation-induced lymphopenia through kinetic sparing of circulating lymphocytes

**DOI:** 10.64898/2026.08.15.744675

**Authors:** Cezara Cheptea, Pierre Loap, Andrew Friberg, Kathryn H. Brown, Ioannis Paraskevaidis, Kristianna Kolker, Michele Kim, Mihaela Ghita-Pettigrew, Mark McDowell, Shiva Shahrampour, Bonnie Ky, Kevin Teo, James Metz, Constantinos Koumenis, Jufri Setianegara, Eric Diffenderfer, Jennifer Wei Zou, Karl T. Butterworth, Ioannis I. Verginadis

## Abstract

**Background and purpose:** Radiation-induced lymphopenia is associated with adverse outcomes in thoracic malignancies. FLASH radiotherapy delivers radiation over a timescale of hundreds of milliseconds, potentially reducing the fraction of irradiated circulating lymphocytes. In this study, we investigated whether FLASH mitigates lymphopenia after thoracic irradiation delivered with protons or photons.

**Materials and methods:** C57BL/6 mice received three 13.5-Gy whole-heart fractions at 48-hour intervals using FLASH or standard dose-rate proton irradiation at the University of Pennsylvania (n=15), with photon validation at Queen’s University Belfast (n=72). Leukocytes and CD4^+^ T cells, CD8^+^ T cells, B cells, and NK cells were quantified by hemocytometer and flow cytometry. A continuous-time Markov model simulated lymphocyte trafficking, dose accumulation, and post-irradiation recovery.

**Results:** FLASH attenuated leukocyte depletion across both proton and photon irradiation modalities. In the proton cohort, white blood cell counts were significantly higher after FLASH at D1, D3, D7, and D14; CD4^+^ T cells and NK cells were preserved through D14, while CD8^+^ T cell sparing persisted through D21. Photon FLASH preserved CD45^+^ leukocytes at D1, D3, D7, and D21, with sustained CD8^+^ sparing at D21. Modeling showed that FLASH shifted the lymphocyte dose distribution toward lower exposures, increasing the proportion of lymphocytes receiving <1 Gy from 2.4% to 16.4%, and reduced the proportion of lymphocytes repeatedly irradiated across all three fractions from 36.3% at standard dose rate to 9.18%, despite similar median cumulative doses. The spleen contributed substantially to cumulative lymphocyte dose, and marrow-entering lymphocytes displayed a more high-dose-enriched distribution after FLASH irradiation.

**Conclusion:** FLASH consistently mitigated radiation-induced lymphopenia for proton and photon modalities, with durable CD8^+^ T cell preservation. These findings support a kinetic mechanism and provide a rationale for combining FLASH radiotherapy with immune-sparing planning and immunotherapy.

## Introduction

Radiation-induced lymphopenia is recognized as an adverse prognostic factor across solid malignancies [1–4]. In thoracic cancers, particularly lung and esophageal carcinomas, severe treatment-related lymphopenia has been independently associated with worse survival [2,3]. This may partly result from irradiation of circulating immune cells as they transit highly perfused organs, including the heart, lungs, and great vessels [5–7]. Accordingly, immune system sparing has emerged as a potential consideration in treatment optimization, supported by dosimetric approaches such as the effective dose to immune cells (EDIC) [8], which estimates radiation exposure to circulating immune cells based on dose delivered to highly perfused organs [9–11]. Because lymphocytes are highly radiosensitive [12,13], with an estimated LD50 of approximately 2 Gy [14], their depletion is influenced by both the volume of irradiated blood and the duration of beam delivery.

FLASH radiotherapy delivers radiation at ultra-high dose rates (UHDR) and has demonstrated reduced normal-tissue toxicity while maintaining antitumor efficacy [15–20]. While the parameters and mechanisms underlying the FLASH effect remain to be fully understood, the markedly shortened beam-on time may provide a specific theoretical mechanism for lymphocyte sparing [21,22]. Conventional deliveries lasting several minutes allow repeated passage of circulating blood through the irradiated field. In contrast, UHDR exposures are completed within hundreds of milliseconds, and may expose a much smaller fraction of the circulating lymphocyte pool and reduce repeated irradiation across fractions.

Previous studies have reported accelerated T cell recovery after UHDR total-body irradiation and reduced immune-cell depletion after whole-thorax irradiation [23,24]. However, these models broadly expose hematopoietic, lymphoid, and circulating compartments and do not reproduce the partial-volume distributions of focal thoracic radiotherapy. Consequently, the longitudinal effects of UHDR exposures on lymphocyte subsets after targeted thoracic irradiation remain uncertain.

Here, we investigated whether UHDR exposures could mitigate radiation-induced lymphopenia after fractionated cardiac irradiation using proton and photon irradiation modalities. We characterized total leukocyte and lymphocyte-subset kinetics and integrated these experimental findings with computational modeling of lymphocyte trafficking and dose accumulation. This combined approach aimed to define the immune-sparing effects of FLASH across radiation modalities and their potential relevance for future immune-preserving treatment strategies.

## Materials and Methods

### Animals

C57BL/6 mice aged 10–11 weeks were used in both institutions. For the proton experiments at the University of Pennsylvania (UPenn, Philadelphia, PA, USA), 15 female mice obtained from The Jackson Laboratory (Bar Harbor, ME, USA) were randomized to non-irradiated controls (NR), FLASH proton irradiation (F-PRT), or standard dose-rate proton irradiation (S-PRT) groups (n = 5 each). For the photon experiments at Queen’s University Belfast (QUB, Belfast, UK), 72 male and female mice obtained from Charles River Laboratories (UK) were equally allocated by sex to NR, FLASH photon irradiation (F-xRT), or conventional dose-rate photon irradiation (S-xRT) groups (n = 24 each). At both institutions, animals were maintained under a 12-hour light/dark cycle with food and water ad libitum and were anesthetized with isoflurane (3% in oxygen at 1 L/min at UPenn and in room air at QUB). UPenn animals were housed in AAALAC-accredited University Laboratory Animal Resources facilities, with procedures approved by the Institutional Animal Care and Use Committee. QUB procedures complied with the Animals (Scientific Procedures) Act 1986 under project license PPL2935 and were approved by the Animal Welfare and Ethical Review Body.

### Irradiation procedure

For proton experiments at UPenn, protons were delivered on the fixed beam line of an IBA Proteus Plus C230 cyclotron (230 MeV; range ≈ 32 g/cm^2^) in a dedicated preclinical room. Field size and flatness were verified before each fraction using Gafchromic EBT3 film. Absolute dosimetry used a NIST-traceable Advanced Markus chamber according to IAEA TRS-398, with online monitoring by a cross-calibrated PTW Bragg Peak transmission chamber. Daily phantom- and-film quality assurance verified imaging-beam isocenter coincidence, and measured shifts were applied during mouse setup. Each mouse was positioned upright in a motorized fixture, receiving isoflurane anesthesia, receiving isoflurane anesthesia. The heart was localized by CBCT and centered using laser guidance and motorized positioning. A double-scattered, posterior shoot-through beam was delivered horizontally towards the heart through an 8-mm brass collimator positioned 1 cm from the skin. Three 13.5-Gy fractions were delivered at 48-hour intervals on D -4, D -2, and D0. A three-fraction regimen was selected (Fig. 1a) to approximate clinically-used stereotactic lung radiotherapy schedules and to provide greater translational relevance. Mean dose rates were 188.76 ± 15.27 Gy/s for F-PRT and 0.74 ± 0.08 Gy/s for S-PRT; beam current was the sole delivery variable. Mean measured dose was 13.52 ± 0.10 Gy/fraction. NR mice underwent identical anesthesia, imaging, and positioning without irradiation.

**Fig. 1.**
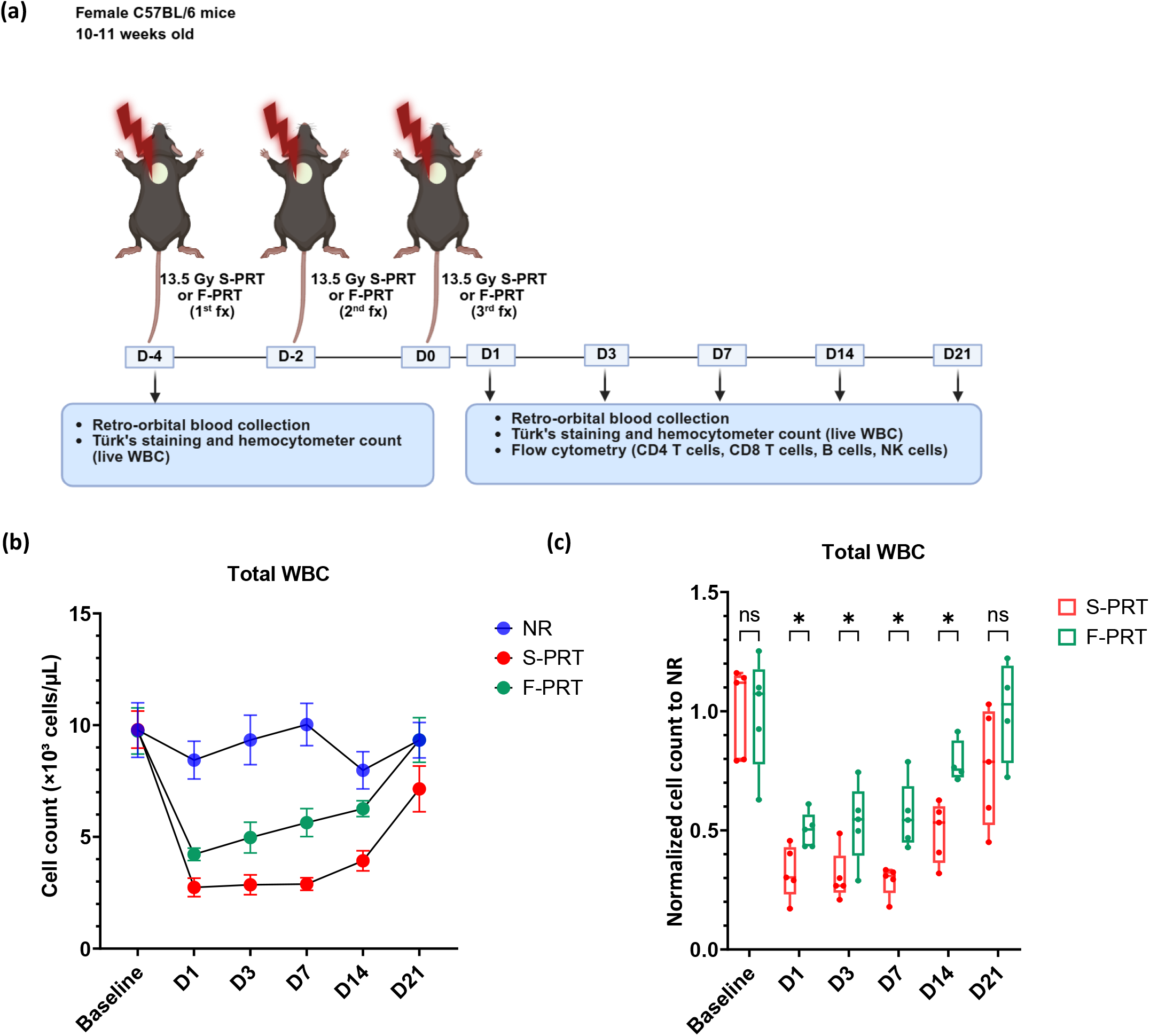
Experimental design and longitudinal changes in white blood cell counts after fractionated proton irradiation. **(a)** Experimental workflow. Mice were assigned to non-irradiated controls (NR, n = 5), standard dose-rate proton irradiation (S-PRT, n = 5), or FLASH proton irradiation (F-PRT, n = 5). Irradiated mice received three fractions of 13.5 Gy delivered at 48-hour intervals. At baseline, corresponding to the day of the first irradiation fraction, retro-orbital blood sampling was performed for white blood cell (WBC) counting using Türk’s solution staining and a hemocytometer. Additional retro-orbital blood samples were collected at day 1, day 3, day 7, day 14, and day 21 after completion of the final irradiation fraction. At each post-irradiation time point, WBC counts were assessed using Türk’s solution staining, and lymphocyte subsets, including CD4^+^ T cells, CD8^+^ T cells, B cells, and NK cells, were analyzed by flow cytometry. **(b)** Absolute live WBC counts measured by Türk’s solution staining and hemocytometer counting over time. Data are presented as mean ± SEM. **(c)** WBC counts in S-PRT and F-PRT mice normalized to the mean value of the NR group at each corresponding time point. Center lines indicate the median, boxes the interquartile range, and whiskers the minimum-to-maximum range. F-PRT is shown in green and S-PRT in red. Statistical comparisons were performed using two-sided unpaired Student’s t-tests. ns, not significant; *, *P* < 0.05. Abbreviations: F-PRT, FLASH proton radiation therapy; S-PRT, standard-dose-rate proton radiation therapy; WBC, white blood cell.

At QUB, photon irradiation used a 150-kVp SARRP-FLASH system with two parallel-opposed X-ray sources, 0.025-mm Cu filtration, 3.4-mm Al HVL, 8-mm collimators, and a 20-mm intertube gap [25]. Heart positioning was established by planar imaging of an anatomical phantom and reproduced using sternal, rib, and vertebral landmarks. Three prescribed 13.5-Gy fractions were delivered on D-4, D-2, and D0. Mean measured dose was 38.68 ± 2.42 Gy in total, or 12.8 ± 0.10 Gy/fraction. F-xRT dose rates were 69.0 ± 2.6 Gy/s at the surface and 60.3 ± 2.26 Gy/s through the heart; corresponding S-xRT rates were 0.68 ± 0.08 and 0.59 ± 0.06 Gy/s. F-xRT was delivered as a continuous UHDR exposure, whereas S-xRT used a pulsed delivery to achieve the lower mean dose rate. EBT4 film was used to monitor dose and field uniformity for each mouse.

### Blood collection, absolute leukocyte counting, and flow cytometry

At UPenn, peripheral blood was collected longitudinally by retro-orbital sampling under isoflurane anesthesia (3% in oxygen, 1 L/min), before the first fraction on D -4 and at D1, D3, D7, D14, and D21 after the final fraction on D0 (Fig. 1a). At QUB, a terminal cross-sectional design was used, with six mice per group euthanized at D1, D3, D7, and D21. At both sites, 50 µL of whole blood was collected into EDTA-coated tubes and maintained on ice; 5 µL was used for absolute leukocyte counting and 45 µL for flow cytometry.

Total leukocytes were quantified using Türk’s solution (Sigma-Aldrich, 1.09277.0100) and a Neubauer hemocytometer. Five microliters of blood were mixed with 95 µL of Türk’s solution, corresponding to a 1:20 dilution, and loaded into the counting chamber. Leukocytes were counted in the four large corner squares, and the mean count was used to calculate the absolute leukocyte concentration per microliter of whole blood.

For flow cytometry, erythrocytes were lysed with 500 µL of ACK buffer (Thermo Fisher Scientific, A1049201) for 3-4 minutes at room temperature. Lysis was stopped with 2 mL of FACS buffer, followed by centrifugation, PBS washing, and viability staining with LIVE/DEAD Fixable Aqua (Invitrogen, L34965). The common panel targeted CD45, CD3, CD4, CD8, B220/CD19, and NK1.1. UPenn antibodies were CD45-Pacific Blue (BioLegend, 157212), CD3–APC/Cy7 (BioLegend, 100222), CD4-FITC (BioLegend, 100406), CD8-PE (BioLegend, 100708), CD45R/B220-PerCP-Cyanine5.5 (Thermo Fisher Scientific, eBioscience, 45-0452-82), and NK1.1-PE/Cy7 (BD Pharmingen, 552878). At QUB, CD4-PerCP/Cy5.5 (BioLegend, 100434) and CD19-FITC (BioLegend, 152404) were used, with the remaining markers unchanged. Apoptotic cells were identified by staining with APC-conjugated Annexin V (BioLegend, 640920) in Annexin V Binding Buffer (BioLegend, 422201). Cells were first gated on forward- and side-scatter characteristics to exclude debris and non-cellular events, followed sequentially by exclusion of dead cells, selection of singlets, and identification of CD45^+^ leukocytes. Within viable singlet CD45^+^ cells, B cells were defined as B220^+^ at UPenn or CD19^+^ at QUB, NK cells as NK1.1^+^, and T cells as CD3^+^. Within the CD3^+^ compartment, CD4^+^ T cells were defined as CD4^+^CD8^−^ and CD8^+^ T cells as CD4^−^CD8^+^. Data were analyzed using FlowJo version 10 (BD Biosciences). Absolute subset counts were calculated by multiplying each subset frequency within viable CD45^+^ cells by the corresponding Türk-derived total leukocyte concentration.

### Mathematical modeling

Murine CBCT images acquired during the proton irradiation experiments were exported to RayStation to reconstruct the three-dimensional dose distribution and generate organ-specific dose-volume histograms (DVHs). Lymphocyte kinetics were adapted from the continuous-time Markov model developed by Beekman et al., including the heart, lungs, liver, spleen, Peyer’s patches, lymph nodes, bone marrow, arterial blood, and venous blood [26]. For most compartments, transit times were stochastic and determined using Gillespie’s algorithm based on physiological transition rates using the probability *P*(Δ*t*_*i*_, *j*) = *k*_*i*→*j*_exp(−Δ*t*_*i*_ ⋅ ∑_*j*_ *k*_*i*→*j*_) based on physiological transition rates, *k*_*i*→*j*_.

Cardiac transit, defined as the time required for circulating blood to pass through the cardiac blood pool, was modeled as a fixed 0.6-second period. This short transit time reflects the rapid murine circulation and high heart rate and was estimated from published physiological values, including a total blood volume of 1.8 mL, approximately 10% of the blood volume contained within the heart, and a cardiac output of approximately 15 mL/min, adjusted for anesthesia [27,28]. For each 13.5-Gy fraction, the irradiation was modeled using a beam-on time of 0.4 seconds for FLASH irradiation and 400 seconds for conventional irradiation.

Dose accumulation was simulated for 100,000 lymphocyte particles (LPs), defined as individual computational agents representing lymphocytes within the circulation model. Each LP was tracked independently as it trafficked between physiological compartments and accumulated radiation dose over time and across fractions. Whenever an LP traversed an irradiated organ, dose was sampled from the corresponding static organ DVH and accumulated over time and across the three fractions. This framework generated cumulative LP DVHs, dose-frequency histograms, and organ-specific contributions to lymphocyte exposure. To investigate potential signaling from irradiated lymphocytes to hematopoietic progenitors, dose-volume statistics for LPs located within the bone marrow were also simulated and data were collected each hour for five days after irradiation.

Lymphocyte death after each fraction was estimated using a linear model with α = 0.3 Gy^−1^ [13]. Post-irradiation population recovery was modeled using three-parameter delay differential equation:

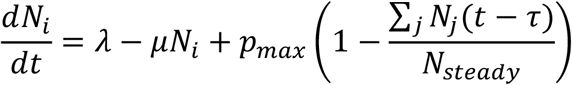

where *N*_*i*_(*t*) denotes the lymphocyte population at time t in compartment i, *N*_*steady*_ the steady-state population, λ the basal production rate, and μ the basal death rate. The parameter *p*_*max*_ represents the maximum inducible excess proliferation rate above basal production, while τ represents the delay between lymphocyte depletion and the resulting compensatory proliferative response. Baseline production was constrained as *λ=μN*_*steady*_ to maintain equilibrium in the absence of irradiation. Model parameters were estimated by parameter sweep to identify the combination providing the best fit to the observed mean longitudinal lymphocyte counts for each subset and irradiation condition.

### Statistical analysis

Absolute leukocyte counts are presented as mean ± standard error of the mean (SEM), unless otherwise specified. Normalized immune-cell counts are shown as box-and-whisker plots, with center lines indicating the median, boxes the interquartile range, and whiskers the minimum-to-maximum range. For normalization, immune-cell counts at each time point were divided by the corresponding mean value of the non-irradiated (NR) control group.

In the UPenn proton cohort, blood samples were collected longitudinally from the same animals, whereas the QUB photon cohort used an independent cross-sectional design at each time point. FLASH and standard dose-rate groups were compared separately at each time point using two-sided unpaired Student’s *t*-tests; no statistical comparisons were performed across time points. All tests were two-sided, with *P* < 0.05 considered statistically significant. Statistical analyses were performed using GraphPad Prism version 10 (GraphPad Software, Boston, MA, USA).

## Results

White blood cell (WBC) counts were compared among NR, S-PRT, and F-PRT groups (Fig. 1b). Following three fractions, both irradiated groups developed rapid leukopenia, with a nadir at D1. However, WBC counts decreased to 2.74 × 10^3^ cells/µL after S-PRT versus 4.22 × 10^3^ cells/µL after F-PRT, corresponding to approximately 1.5-fold higher counts in the FLASH group, while remaining stable in NR mice. After normalization to the NR mean at each time point, WBC counts were significantly higher after F-PRT than S-PRT at D1, D3, D7, and D14 (all P < 0.05; Fig. 1c). By D21, both groups had partially recovered and no longer differed significantly. Flow-cytometric analysis revealed subset-specific lymphocyte sparing after F-PRT (Fig. 2). Compared with S-PRT, CD4^+^ T cell counts were significantly higher at D1, D3, D7, and D14, before converging by D21 (Fig. 2a). CD8^+^ T cell counts were significantly higher after F-PRT at every time point from D1 through D21 (Fig. 2b). B cells were markedly depleted in both irradiated groups, with no significant difference at D3 or D21 (Fig. 2c). NK-cell counts were significantly higher after F-PRT at D1, D3, D7, and D14 (Fig. 2d). Annexin V positivity was comparable between F-PRT and S-PRT across lymphocyte subsets and time points, except for NK cells at D21, which showed significantly higher Annexin V positivity following S-PRT than F-PRT (Supplementary Fig. 1). At D21, Annexin V positivity was also numerically increased relative to earlier time points in both B and NK cells in the two irradiated groups.

**Fig. 2.**
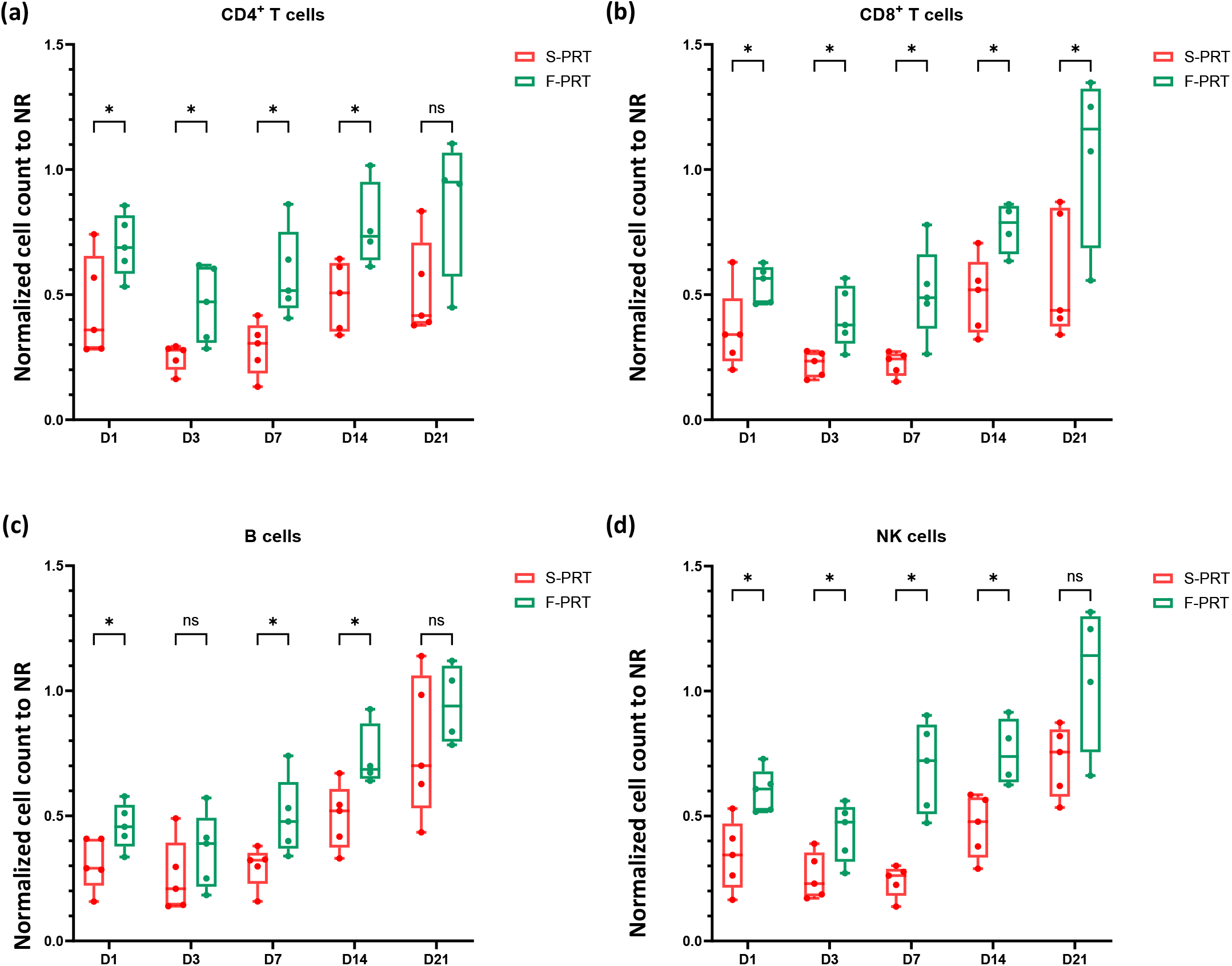
Longitudinal changes in circulating immune cell counts after fractionated proton irradiation. Circulating counts of CD4^+^ T cells. **(a)**, CD8^+^ T cells **(b)**, B cells **(c)**, and NK cells **(d)**, normalized to the mean value of the non-irradiated control group, in mice treated with FLASH proton irradiation (F-PRT, n = 5) or standard dose-rate proton irradiation (S-PRT, n = 5). Mice received three fractions of 13.5 Gy delivered at 48-hour intervals. Immune cell counts were measured at 1, 3, 7, 14, and 21 days after completion of the final irradiation fraction. For all box- and-whisker plots, center lines indicate the median, boxes the interquartile range, and whiskers the minimum-to-maximum range. F-PRT is shown in green and S-PRT in red. Statistical comparisons were performed using two-sided unpaired Student’s t-tests. ns, not significant; *, *P* < 0.05. Abbreviations: F-PRT, FLASH proton radiation therapy; S-PRT, standard-dose-rate proton radiation therapy.

To assess whether FLASH-mediated leukocyte sparing was reproducible across radiation modalities, we analyzed an independent photon irradiation cohort at Queen’s University Belfast. Both S-xRT and F-xRT groups developed an early decline in circulating CD45^+^ leukocytes, with a nadir at D1 followed by progressive recovery. At D1, mean CD45^+^ leukocyte counts were 3.23 × 10^3^ cells/µL after S-xRT and 8.03 × 10^3^ cells/µL after F-xRT, corresponding to an approximately 2.5-fold difference. CD45^+^ leukocyte counts remained significantly higher after F-xRT than after S-xRT at D1, D3, D7, and D21 (Fig. 3a-b). Analysis of lymphocyte subsets showed an overall trend toward greater preservation with FLASH, although statistical significance varied by subset and time point (Fig. 3c-f). B cell counts were significantly higher at D1, D3, and D21. Preservation of CD4^+^ T cell counts were significantly higher after F-xRT at D1 and tended to remain numerically higher thereafter. CD8^+^ T cell counts were significantly higher at D1 and D21, with a similar trend at D3 and D7. NK-cell preservation was significant at D1 only. These findings independently confirm the leukocyte-sparing effect of FLASH and indicate that it is not restricted to proton thoracic irradiation.

**Fig. 3.**
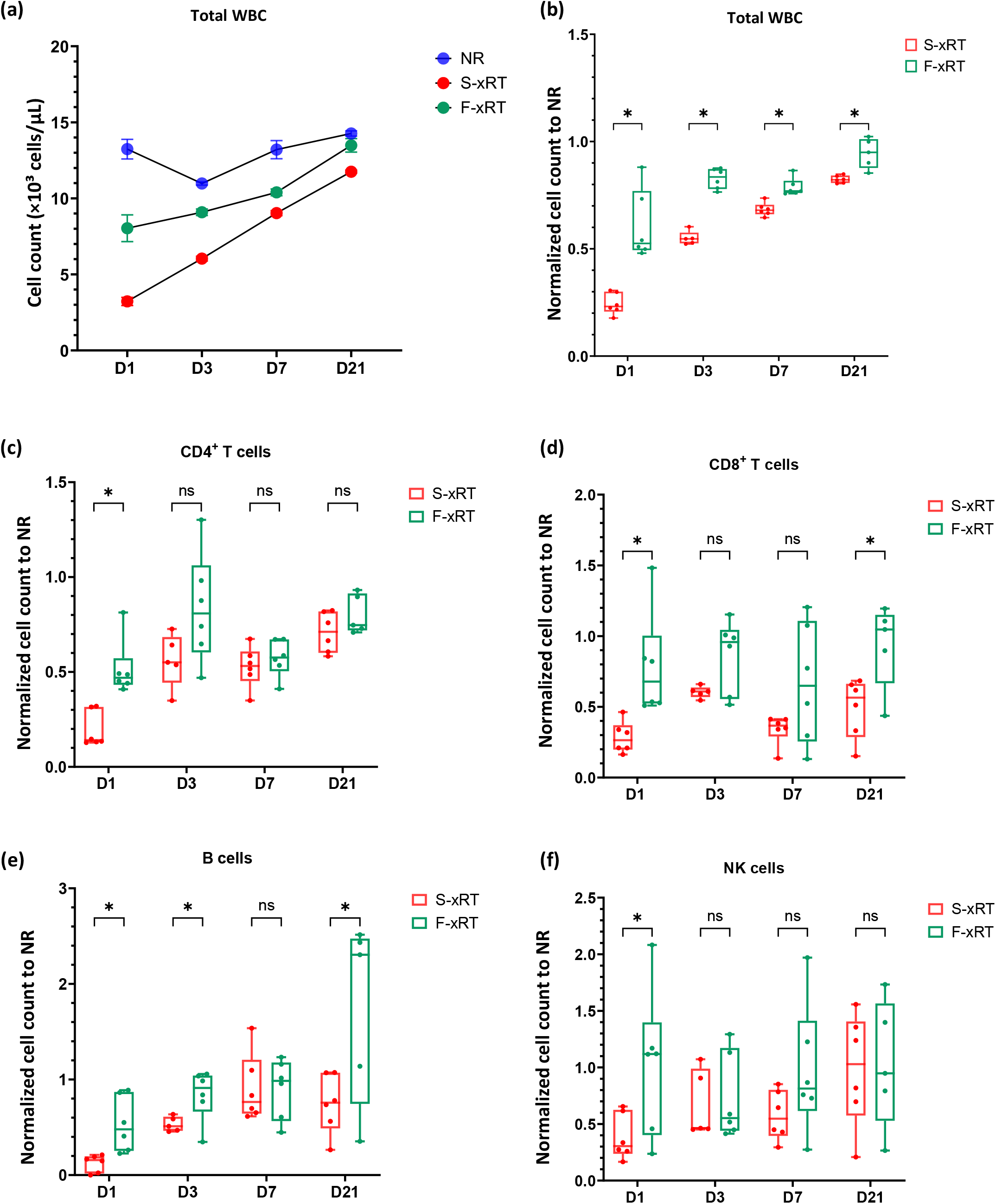
Independent validation of lymphocyte sparing after FLASH photon irradiation. Circulating white blood cells (WBC) and lymphocyte subsets were quantified after fractionated FLASH (F-xRT) or standard dose-rate (S-xRT) photon irradiation at Queen’s University Belfast. **(a)** Absolute WBC counts. Data are presented as mean ± SEM. **(b)** WBC counts after F-xRT and S-xRT, normalized to the mean of the corresponding non-irradiated control group (NR) at each time point. **(c–f)** CD4^+^ T cell, CD8^+^ T cell, B cell, and NK-cell counts, respectively, after F-xRT and S-xRT, normalized to the corresponding NR mean. Measurements were performed at D1, D3, D7, and D21 after the final irradiation. For all box-and-whisker plots, center lines indicate the median, boxes the interquartile range, and whiskers the minimum-to-maximum range. F-xRT is shown in green and S-xRT in red. Statistical comparisons were performed using two-sided unpaired Student’s t-tests. ns, not significant; *, *P* < 0.05. Abbreviations: F-xRT, FLASH X-ray radiation therapy; S-xRT, standard-dose-rate X-ray radiation therapy; WBC, white blood cell.

Predicted cumulative dose distributions for circulating lymphocytes differed substantially between FLASH and standard dose-rate proton irradiation (Fig. 4a), despite similar median cumulative doses of 9.32 and 9.66 Gy, respectively. Simulations showed that FLASH shifted the modeled lymphocyte dose distribution toward lower exposures, increasing the proportion of lymphocyte particles receiving less than 1 Gy from 2.4% to 16.4% and reducing the proportion irradiated during all three fractions from 36.3% to 9.18% (Fig. 4b-d). Thus, FLASH reduced repeated irradiation across fractions while increasing the proportion of lymphocytes remaining within the minimally exposed dose range. The model further predicted that cumulative lymphocyte dose was not determined solely by transit through the irradiated heart. Although the heart and lungs received the highest static organ doses, the spleen was predicted to contribute the largest share of cumulative lymphocyte exposure, whereas cardiac transit contributed comparatively little (Fig. 4e-f).

**Fig. 4.**
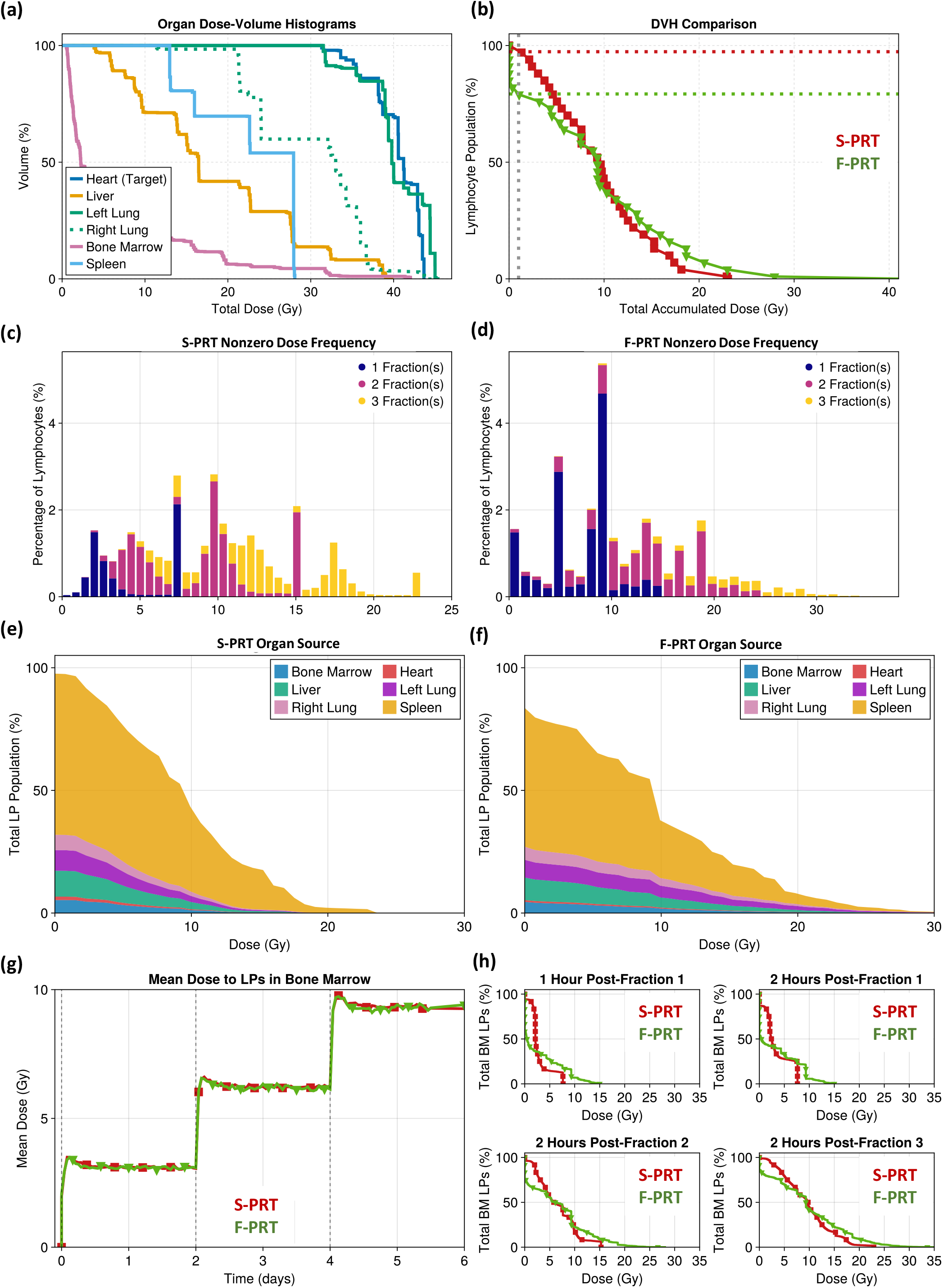
Modeling of lymphocyte dose accumulation during fractionated cardiac proton irradiation. **(a)** Static dose-volume histograms (DVHs) of the irradiated organs. **(b)** Cumulative dose–volume histograms of simulated lymphocyte particles (LPs) after completion of the three treatment fractions; dotted lines indicate the 1 Gy threshold. **(c, d)** Dose-frequency histograms showing the number of fractions during which each LP was irradiated under standard dose-rate (S-PRT, c) and FLASH (F-PRT, d) conditions. **(e, f)** Organ-specific breakdown of cumulative LP dose, showing the contribution of each compartment to total lymphocyte exposure under standard dose-rate (S-PRT, e) and FLASH (F-PRT, f) irradiation. **(g)** Predicted mean dose of LPs present within the bone marrow compartment during and after treatment. **(h)** DVHs of LPs within the bone marrow compartment shortly after each fraction. LP, lymphocyte particle. Abbreviations: F-PRT, FLASH proton radiation therapy; S-PRT, standard-dose-rate proton radiation therapy.

Irradiated lymphocytes subsequently reached the bone marrow within hours, where their dose distribution rapidly stabilized. Mean doses in bone-marrow-resident particles were similar across dose-rate conditions, but FLASH produced a more heterogeneous DVH, combining a smaller low-dose population with a more pronounced high-dose tail (Fig. 4g-h). Fitting the delay-differential model to the experimentally observed longitudinal lymphocyte counts required a non-zero signaling delay between radiation-induced depletion and the onset of compensatory proliferation. This delay was longer after standard dose-rate irradiation for CD4^+^ T cells, CD8^+^ T cells, and NK cells, consistent with a slower recovery (Supplementary Fig. 2).

Overall, these simulations predict that FLASH alters the lymphocyte dose distribution rather than uniformly reducing dose, preserving a larger fraction of minimally exposed circulating cells and limiting repeated irradiation across fractions, while producing a more high-dose-enriched DVH among lymphocytes subsequently reaching the bone marrow.

## Discussion

This study shows that UHDR irradiation mitigates radiation-induced lymphopenia after focal thoracic irradiation. Fractionated cardiac irradiation induced a rapid decline in circulating leukocytes, with the greatest depletion observed at D1 after conventional dose-rate irradiation, while this decrease was attenuated under FLASH conditions. This effect was reproduced with proton irradiation at the University of Pennsylvania and photon irradiation at Queen’s University Belfast despite differences in irradiation modality, geometry, and experimental design, supporting a dose-rate-dependent response that is consistent across both proton and photon irradiation.

The most consistent subset-level finding across both proton and photon modalities was preservation of CD8^+^ T cells. In the proton cohort, CD8^+^ T cell counts remained significantly higher after FLASH throughout follow-up, whereas the photon cohort showed significant sparing at D1 and D21, suggesting that FLASH may reduce both the magnitude and persistence of CD8^+^ T cell depletion. Other lymphocyte subsets showed less consistent patterns across experiments. CD4^+^ T cell and NK-cell preservation was sustained through D14 in the proton cohort, whereas in the photon cohort CD4^+^ T cell sparing reached significance only at D1 and NK-cell sparing only at D1. Conversely, B cell sparing was limited in the proton cohort but was significant after photon FLASH at D1, D3, and D21. These differences should not necessarily be interpreted as modality-specific biological effects, as the two experiments also differed in FLASH dose rate, blood-sampling design (serial versus terminal sampling), and sex composition of the cohorts. Sustained

CD8^+^ T cell preservation may nevertheless be clinically relevant, given the central role of CD8^+^ T cells in antitumor immunity and immune-checkpoint inhibition [29]. This may be particularly pertinent in thoracic and breast malignancies, where radiotherapy is commonly combined with durvalumab in locally advanced lung cancer [7,30], nivolumab in selected esophageal or gastroesophageal junction cancers [31], and pembrolizumab in triple-negative breast cancer [32]. However, preservation of circulating CD8^+^ T cell counts does not necessarily imply preserved effector function or increased tumor infiltration. Tumor-associated inflammation may alter lymphocyte kinetics and trafficking after irradiation [33], while immune-checkpoint inhibition may further modify T cell activation, proliferation, and recruitment to the tumor microenvironment [34]. Whether FLASH-mediated preservation of circulating lymphocytes ultimately enhances intratumoral immune responses or the efficacy of immune-checkpoint inhibitors therefore remains to be established in tumor-bearing models.

Our findings align with several preclinical studies reporting lymphocyte-sparing effects of FLASH irradiation despite the use of relatively large treatment fields. Tao et al. observed attenuated lymphopenia and more rapid hematologic recovery after whole-thorax FLASH irradiation [24], consistent with the recovery pattern observed in our models. Our study extends these findings by demonstrating concordant immune-cell sparing after partial thoracic irradiation across both proton and photon modalities. Using total-body irradiation, Yu et al. found no major difference in peripheral blood counts between FLASH and conventional dose-rate delivery but reported faster T cell recovery after FLASH, supporting an effect on post-irradiation lymphocyte kinetics [23]. However, compartment-specific responses may nevertheless complicate this interpretation; Qin et al. described a dual effect in which FLASH irradiation of limb preserved circulating lymphocytes while exacerbating lymphocyte depletion in the non-irradiated spleen [35].

Computational studies have also reached differing conclusions. Our model, adapted from the continuous-time Markov framework developed by Beekman et al. [26], supports a kinetic mechanism in which shorter beam-on time limits both the fraction of circulating lymphocytes exposed and their repeated irradiation across fractions. In contrast, the LymphoDose framework developed by de Kermenguy et al. [36], when applied to a glioma cohort, predicted only modest differences between ultra-high and conventional dose-rate irradiation and attributed most lymphocyte exposure to lymph nodes rather than circulating blood. Together, these findings suggest that FLASH-mediated immune sparing depends strongly on treatment site, field geometry, and the relative contributions of circulating and tissue-resident lymphocyte compartments.

In our thoracic irradiation model, the experimental and computational findings support a predominantly kinetic mechanism of lymphocyte sparing. By markedly shortening beam-on time, FLASH reduces the fraction of circulating blood exposed during irradiation and the likelihood of repeated lymphocyte exposure across fractions. Accordingly, the model predicted a redistribution of the lymphocyte DVH toward lower-dose exposure, increasing the proportion of lymphocytes receiving less than 1 Gy from 2.4% to 16.4%, with a corresponding reduction in the proportion receiving higher doses, and a reduction in the proportion irradiated during all three fractions from 36.3% to 9.18%, despite similar median cumulative doses. Given the high radiosensitivity of lymphocytes [13], this redistribution toward lower and less repetitive exposure is likely biologically meaningful. Consistent with this interpretation, Annexin V positivity was largely comparable between dose-rate groups, except for higher NK-cell Annexin V positivity after S-PRT at D21, suggesting that the overall lymphocyte-sparing effect of FLASH primarily resulted from reduced exposure rather than a broadly differential apoptotic response.

The delay-differential model also indicated faster recovery after FLASH and a distinct dose distribution among lymphocyte particles reaching the bone marrow. Although FLASH preserved a larger minimally exposed population overall, marrow-entering lymphocytes showed a higher-dose-enriched DVH, with more heavily irradiated cells and fewer cells within the low-dose range. Because irradiated lymphocytes traffic through the marrow [37], this difference could potentially alter compensatory signaling to hematopoietic progenitors. A smaller but more severely damaged returning population might generate an earlier regenerative signal than the broader, lower-dose exposure produced by standard delivery, although this hypothesis requires experimental validation.

The modeling of lymphocyte trafficking and dose accumulation further showed that immune-cell dose cannot be inferred from cardiac dose alone. Although irradiation was centered on the heart, substantial lymphocyte exposure occurred during transit through other irradiated compartments, particularly the lungs and spleen, with an additional contribution from the liver. These findings have direct implications for immune-sparing treatment planning. The effective dose to immune cells (EDIC) framework integrates dose contributions from highly perfused organs and other exposed compartments, including the heart, lungs, liver, and remaining body [5]. Higher EDIC values have been associated with more severe lymphopenia and poorer outcomes in thoracic malignancies [5,7,38,39], supporting treatment-planning strategies that reduce dose across these structures simultaneously. Proton therapy and FLASH may provide complementary immune-sparing effects: proton therapy reduces integral and low-dose exposure to perfused tissues, whereas FLASH limits the proportion of circulating lymphocytes exposed during beam delivery. Their combination could therefore reduce immune-cell irradiation both spatially and temporally, particularly in thoracic and breast cancers increasingly treated with immunotherapy.

Several limitations should be acknowledged. The proton cohort included only five mice per group, although its findings were supported by the larger photon cohort. Direct comparisons between the two experiments are limited by inter-site differences in irradiation setup and experimental design, including serial blood sampling of the same animals in the proton cohort versus terminal sampling of different animals at each time point in the photon cohort, as well as the use of female mice only at UPenn versus a mixed-sex population at QUB. Both experiments used healthy mice, and tumor-associated inflammation, immunosuppression, and systemic therapies may alter lymphocyte kinetics. Only one fractionation schedule was evaluated, and tumor control was not assessed. Nevertheless, the concordant findings across proton and photon irradiation, together with consistent CD8^+^ sparing and supporting computational modeling, identify immune-cell preservation as a potentially clinically relevant consequence of FLASH delivery. Future studies should prioritize tumor-bearing models, combinations with immune-checkpoint inhibitors, and integration of immune-dose metrics into proton and FLASH treatment planning.

In conclusion, proton and photon UHDR exposures consistently reduced radiation-induced lymphopenia, with sustained preservation of CD8^+^ T cells. These findings provide a strong rationale for evaluating UHDR exposures, particularly proton FLASH irradiation, as an immune-sparing strategy in combination with immunotherapy.

## CRediT authorship contribution statement

**Cezara Cheptea**: Methodology; Investigation; Formal analysis; Data curation; Visualization; Writing – original draft; Writing – review & editing. **Pierre Loap**: Conceptualization; Methodology; Investigation; Formal analysis; Data curation; Visualization; Writing – original draft; Writing – review & editing. **Andrew Friberg**: Methodology; Software; Formal analysis; Data curation; Visualization; Writing – original draft (mathematical modeling); Writing – review & editing. **Kathryn H. Brown**: Methodology; Investigation; Formal analysis; Data curation; Visualization; Writing – original draft (Queen’s University Belfast experimental procedures); Writing – review & editing. **Ioannis Paraskevaidis**: Methodology; Investigation; Formal analysis; Data curation. **Kristianna Kolker**: Investigation; Project administration. **Michele Kim**: Methodology; Investigation; Resources; Writing – original draft (irradiation and dosimetry methods); Writing – review & editing. **Mihaela Ghita-Pettigrew**: Investigation; Formal analysis; Data curation; Writing – original draft (Queen’s University Belfast experimental procedures); Writing – review & editing. **Mark McDowell**: Investigation; Formal analysis; Data curation; Writing – original draft (Queen’s University Belfast experimental procedures); Writing – review & editing. **Shiva Shahrampour**: Methodology; Investigation; Resources. **Bonnie Ky**: Methodology; Investigation. **Kevin Teo**: Methodology; Investigation; Resources. **James Metz**: Resources; Supervision. **Constantinos Koumenis**: Conceptualization; Resources; Supervision; Funding acquisition. **Jufri Setianegara**: Methodology; Investigation; Resources; Writing – original draft (irradiation and dosimetry methods); Writing – review & editing. **Eric Diffenderfer**: Methodology; Investigation; Resources; Supervision; Writing – original draft (irradiation and dosimetry methods); Writing – review & editing. **Jennifer Wei Zou**: Methodology; Supervision; Project administration; Writing – review & editing. **Karl T. Butterworth**: Conceptualization; Methodology; Resources; Supervision; Project administration; Writing – original draft (Queen’s University Belfast experimental procedures); Writing – review & editing. **Ioannis I. Verginadis**: Conceptualization; Methodology; Resources; Supervision; Project administration; Funding acquisition; Writing – review & editing.

## Acknowledgments

The irradiations were performed by the Cell and Animal Radiation Core Facility (RRID:SCR_022377) at the University of Pennsylvania Perelman School of Medicine. We would like to thank the in-house Ion Beam Applications (IBA) physics team for facilitating the preclinical experiments.

## Declaration of generative AI and AI-assisted technologies in the manuscript preparation process

During the preparation of this work the authors used Claude (Anthropic) in order to check the manuscript for language, clarity, and internal consistency, and to edit the text accordingly. After using this tool, the authors reviewed and edited the content as needed and take full responsibility for the content of the published article.

