## Supplementary information for "Fractionated proton and photon FLASH irradiation mitigates radiation-induced lymphopenia through kinetic sparing of circulating lymphocytes"

### Supplementary Figure 1

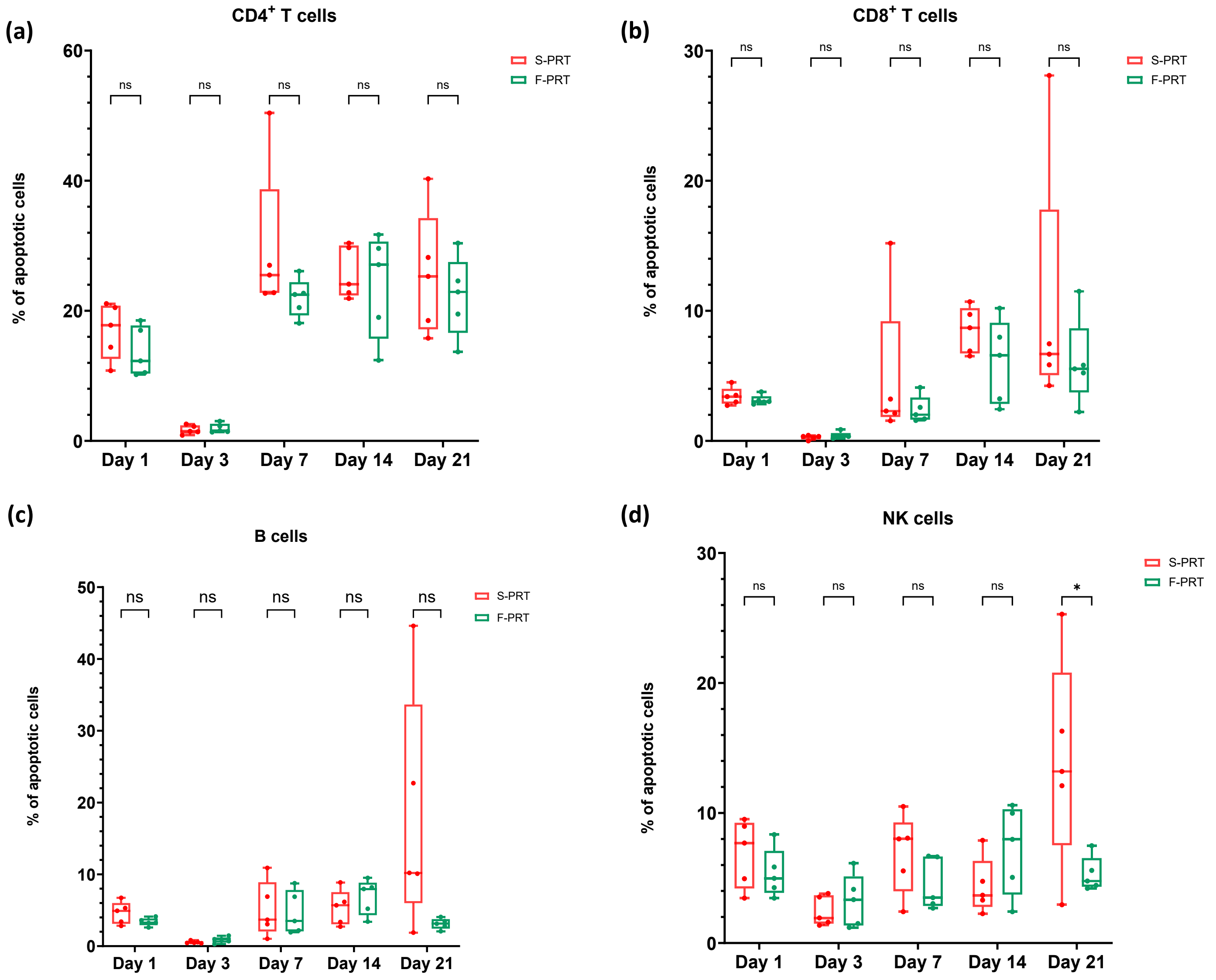

#### Supplementary Fig. 2

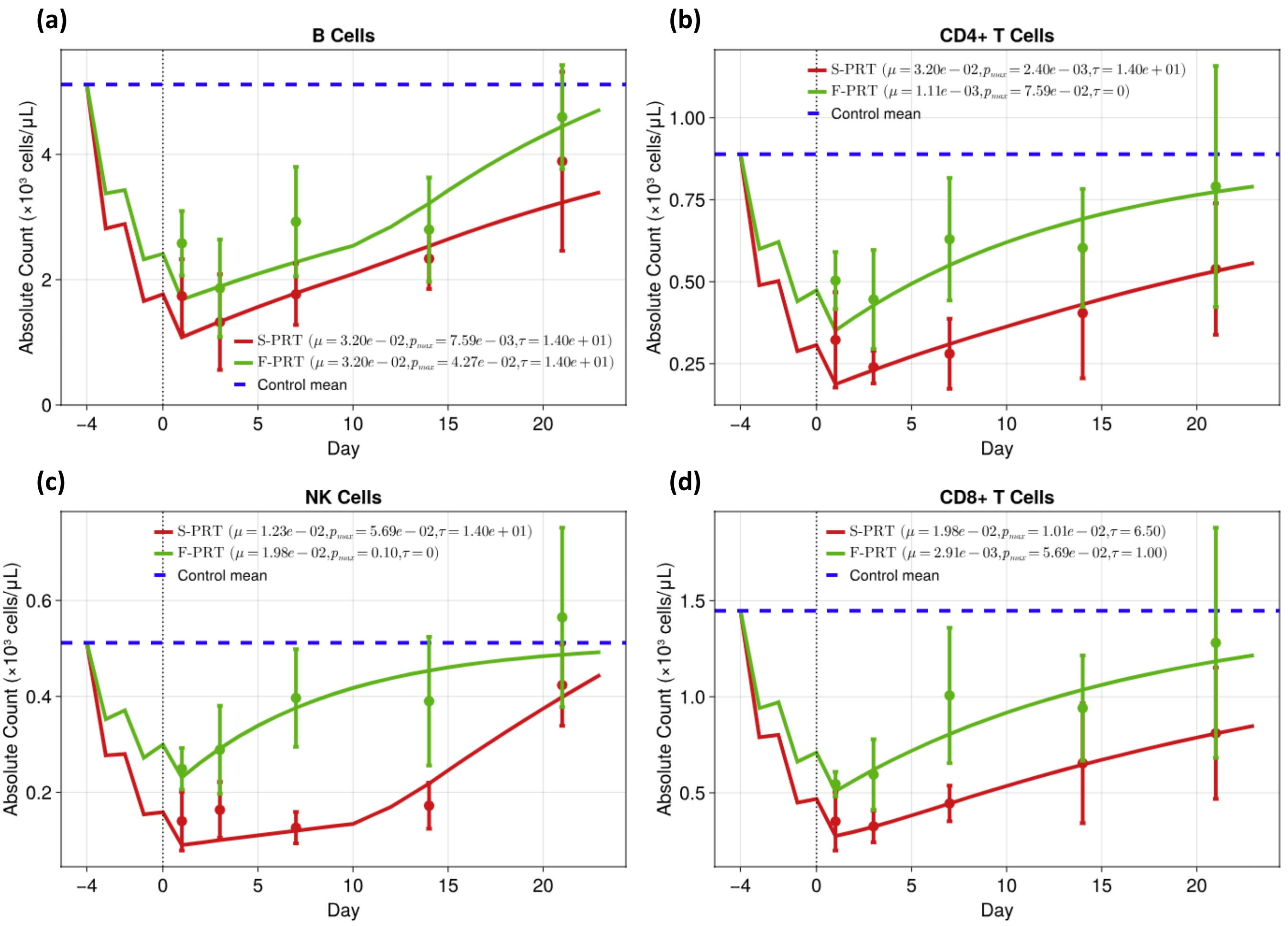

**Supplementary Figure Captions**

**Supplementary Fig. 1. Annexin V positivity in circulating lymphocyte subsets after fractionated proton irradiation.** Proportion of Annexin V+ cells among $CD4^+$ T cells **(a)**, $CD8^+$ T cells **(b)**, B cells **(c)**, and NK cells **(d)**, in F-PRT (n = 5) and S-PRT (n = 5) mice. For all box-and-whisker plots, center lines indicate the median, boxes the interquartile range, and whiskers the minimum-to-maximum range. F-PRT is shown in green and S-PRT in red. Statistical comparisons were performed using two-sided unpaired Student's t-tests. ns, not significant; *, $P < 0.05$. Abbreviations: F-PRT, FLASH proton radiation therapy; S-PRT, standard-dose-rate proton radiation therapy.

**Supplementary Fig. 2. Lymphocyte recovery measurements and DDE model fits.** Population measurements of B cells **(a)**, $CD4^+$ T cells **(b)**, NK cells **(c)**, and $CD8^+$ T cells **(d)** at 1, 3, 7, 14, and 21 days after proton irradiation (D0 corresponding to the third fraction). Recovery modeling curves are shown with basal death rate ($\mu$) and maximum proliferation rate ($p_{max}$) given in days$^{-1}$. Abbreviations: F-PRT, FLASH proton radiation therapy; S-PRT, standard-dose-rate proton radiation therapy.
